# Chronotate: An open-source tool for manual timestamping and quantification of animal behavior

**DOI:** 10.1101/2023.05.31.543063

**Authors:** Paul A. Philipsberg, Zoé Christenson Wick, Keziah S. Diego, Nick Vaughan, Angelina Galas, Albert Jurkowski, Yu Feng, Lauren M. Vetere, Lingxuan Chen, Denise J. Cai, Tristan Shuman

## Abstract

A core necessity to behavioral neuroscience research is the ability to accurately measure performance on behavioral assays, such as the novel object location and novel object recognition tasks. These tasks are widely used in neuroscience research and measure a rodent’s instinct for investigating novel features as a proxy to test their memory of a previous experience. Automated tools for scoring behavioral videos can be cost prohibitive and often have difficulty distinguishing between active investigation of an object and simply being in close proximity to an object. As such, many experimenters continue to rely on hand scoring interactions using stopwatches which makes it difficult to review scoring after-the-fact and results in the loss of temporal information. Here, we introduce Chronotate, a free, open-source tool to aid in manually scoring novel object behavior videos. The software consists of an interactive video player with keyboard integration for marking timestamps of behavioral events during video playback, making it simple to quickly score and review bouts of rodent-object interaction. In addition, Chronotate outputs detailed interaction bout data, allowing for nuanced behavioral performance analyses. Using this detailed temporal information, we demonstrate that novel object location performance peaks within the first 3 seconds of interaction time and preference for the novel location becomes reduced across the test session. Thus, Chronotate can be used to determine the temporal structure of interactions on this task and can provide new insight into the memory processes that drive this behavior. Chronotate is available for download at: https://github.com/ShumanLab/Chronotate.

## INTRODUCTION

The accurate and reliable assessment of behavioral performance in rodents has been of great benefit to virtually every field of neuroscience research from both a basic research and preclinical perspective [1]. Two of the most commonly used rodent behavioral assays are the novel object recognition (NOR) and novel object location (NOL) tasks [2,3]. These two assays test a rodent’s ability to learn the layout and features of an environment and identify changes to that environment after a delay [2]. As such, these tasks test both learning and memory, as a rodent must first learn about an environment and then later remember it well enough to determine which object has changed location or identity. Because rodents have an innate tendency to investigate novel over familiar stimuli [4], memory performance can be measured as a ratio of how much time the rodent spends investigating novel versus familiar stimuli or a new location of familiar stimuli. In fact, this instinct towards investigating novel stimuli or location is so robust that if an equivalent amount of time is spent investigating a novel and a familiar stimulus/location, it is interpreted as the rodent forgetting which is novel and which is familiar [3]. It is therefore critical to the interpretation of this data that the time the rodents spend investigating both novel and familiar stimuli or location be accurately assessed and reported.

A number of machine-learning based tools for automated behavior scoring exist [5–8], but these tools often require specialized video recording setups, large sets of training data, and significant time to fine tune the underlying models [9]. Moreover, subtle distinctions between nuanced behaviors often require the judgment of the experimenter to resolve ambiguous cases. For example, the difference between an animal sniffing an object and an animal grooming while facing an object can be difficult for an automated scoring tool to accurately classify. For these reasons, many experimenters performing the NOR and NOL tasks continue to rely on hand-scoring methods to measure rodent-object investigation times and ratios.

At its simplest, NOR/NOL video scoring requires recording the total time spent interacting with each object. As such, a common method of scoring would consist of using two stopwatches to time the total duration of all bouts of exploration for each of the novel and familiar objects as the video is played back in real-time. This approach has several disadvantages. Recording multiple short bouts of interaction in succession is hindered by the difficulty of pressing the start/stop button of a stopwatch repeatedly, and making even one error can require the scorer to restart from the beginning of the video. Moreover, recording only the total amount of time spent with each object prevents more granular analysis of how interaction changes over the course of the session. If the experimenter is later interested in reexamining only the first 30 seconds of exploration for example, then each video would need to be rescored. Additionally, this approach makes evaluating inter-rater reliability difficult, as scoring of individual bouts cannot be directly compared. Fortunately, however, many of these issues can be addressed by manually noting the duration and timestamp of each bout of interaction, though doing this in a purely manual fashion would be arduous and time-consuming.

Several behavioral scoring software packages support using keystrokes to mark timestamped events on behavioral videos, but the price of these tools can be prohibitive in many applications [2,8]. For this reason, and those discussed above, we developed Chronotate: a free, open-source tool to aid in the hand-scoring of behavior videos. Here we demonstrate Chronotate’s utility for assessing mouse performance on the NOL task and explore object investigation bout-by-bout in addition to surveying cumulative investigation preferences over the duration of testing. We find that not only is Chronotate useful for scoring videos, but that the more granular data analyses it permits can shed light on NOR and NOL behavioral assay design.

## MATERIALS AND METHODS

### Animals

The use of animals and the experimental protocols were approved by the Icahn School of Medicine’s Institutional Animal Care and Use Committee, in accordance with the US National Institutes of Health guidelines.

Male and female C57BL6 mice (Charles River Lab, strain 027, **Fig. 2A**) were ordered directly from the vendor and were tested in the NOL task at 12 weeks of age. In addition, male and female wild-type B6129SF2/J (Jackson Laboratory, RRID: IMSR_JAX: 101045; [10]) and 3xTg (Mouse Mutant Resource and Research Center, RRID:MMRRC_034830-JAX; [11]) were maintained with in-house breeding in approved breeding facilities at Mount Sinai and were tested in the NOL at 6-8 months of age (**Fig. 2B-D, Fig. 3**). All animals were housed in a temperature-controlled vivarium and were given food and water *ad libitum*. Lights were kept on a 12-hour light-dark cycle (lights on at 0700).

### Novel Object Location Memory Task

For 5 days prior to being introduced to the behavioral arena, mice were habituated to transportation from the animal facility to behavior room where they were then gently handled for 5 minutes (**Fig 1A**) [12]. On the 6th and 7th days, mice were habituated to the behavioral arena (30 × 30 × 30cm, white acrylic) for 5 minutes per day. On these days, the arena was empty except for distinct spatial cues on each of the four walls. On the 8th day, mice were returned to the same arena for one (**Fig. 2A, Fig. 3**) to three (**Fig. 2C-D**) training sessions, where two identical objects (plastic toys or door stoppers ∼2” x 2”) were taped down near two corners of the arena (**Fig 1A**). Mice were then returned to their home cage for 1 (**Fig. 2A, Fig. 3**) or 4 (**Fig. 2C-D**) hours before the testing session. During testing, one of the two objects was relocated to a corner of the arena that was previously empty (**Fig 1A**).

**Figure 1:**
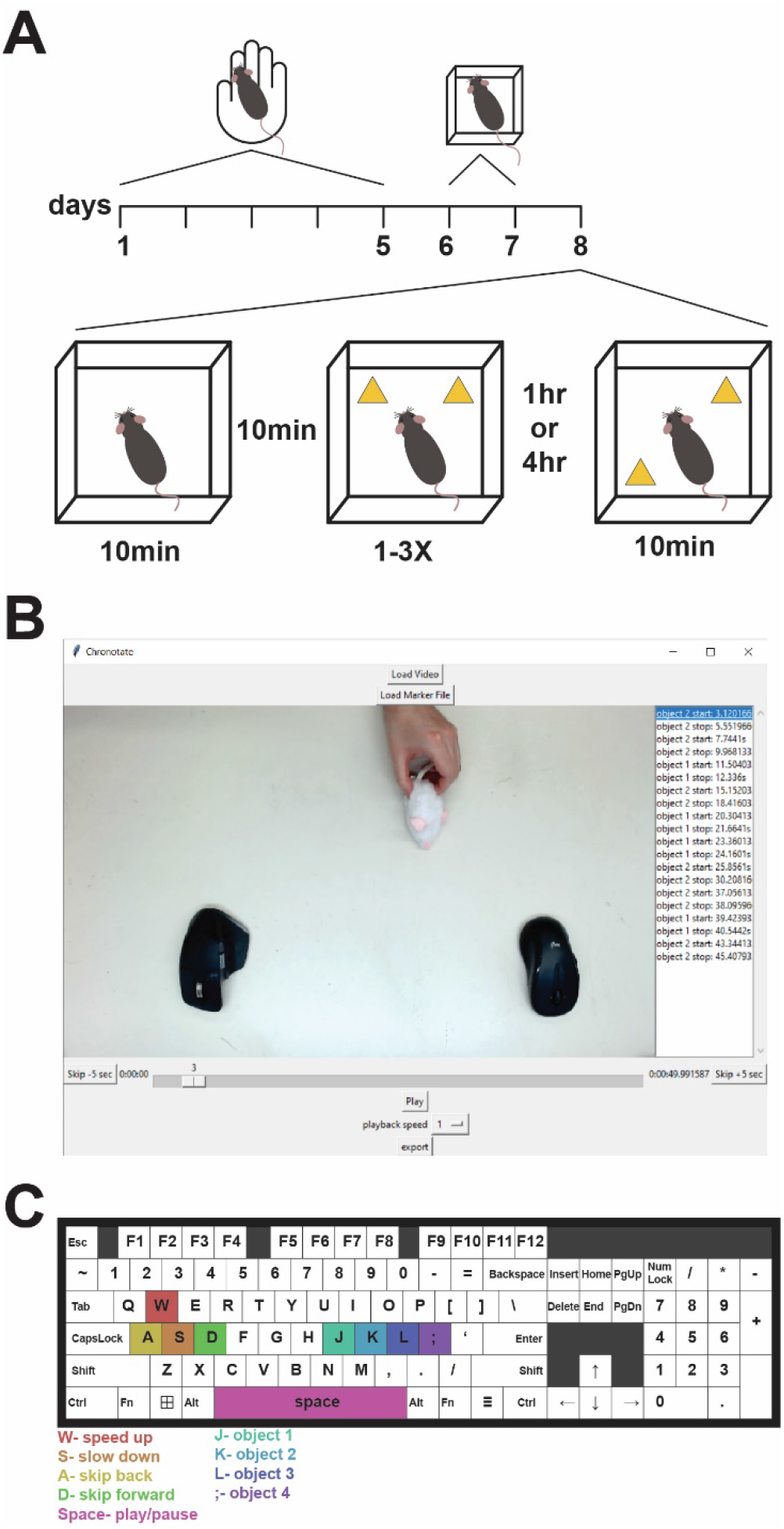
Chronotate for novel object location test scoring. Schematic of novel object location (NOL) protocol. Mice were handled for 5 days before being habituated to an empty arena for 2 days. On the 8^th^ day of training, mice were re-habituated to the empty arena before a training session, where 2 objects were added to the arena. 4 hours following the training session, mice were placed back into the arena with one of the objects relocated. (B) Chronotate graphical interface. A list of time-stamped interaction bouts is listed on the right and a seek/scrub bar along the bottom of the interface. (C) Chronotate keyboard controls for scoring objection interaction (J,K,L,;), speeding up or slowing down (W,S), skipping forward or back (D,A), and playing or pausing (space).

**Figure 2:**
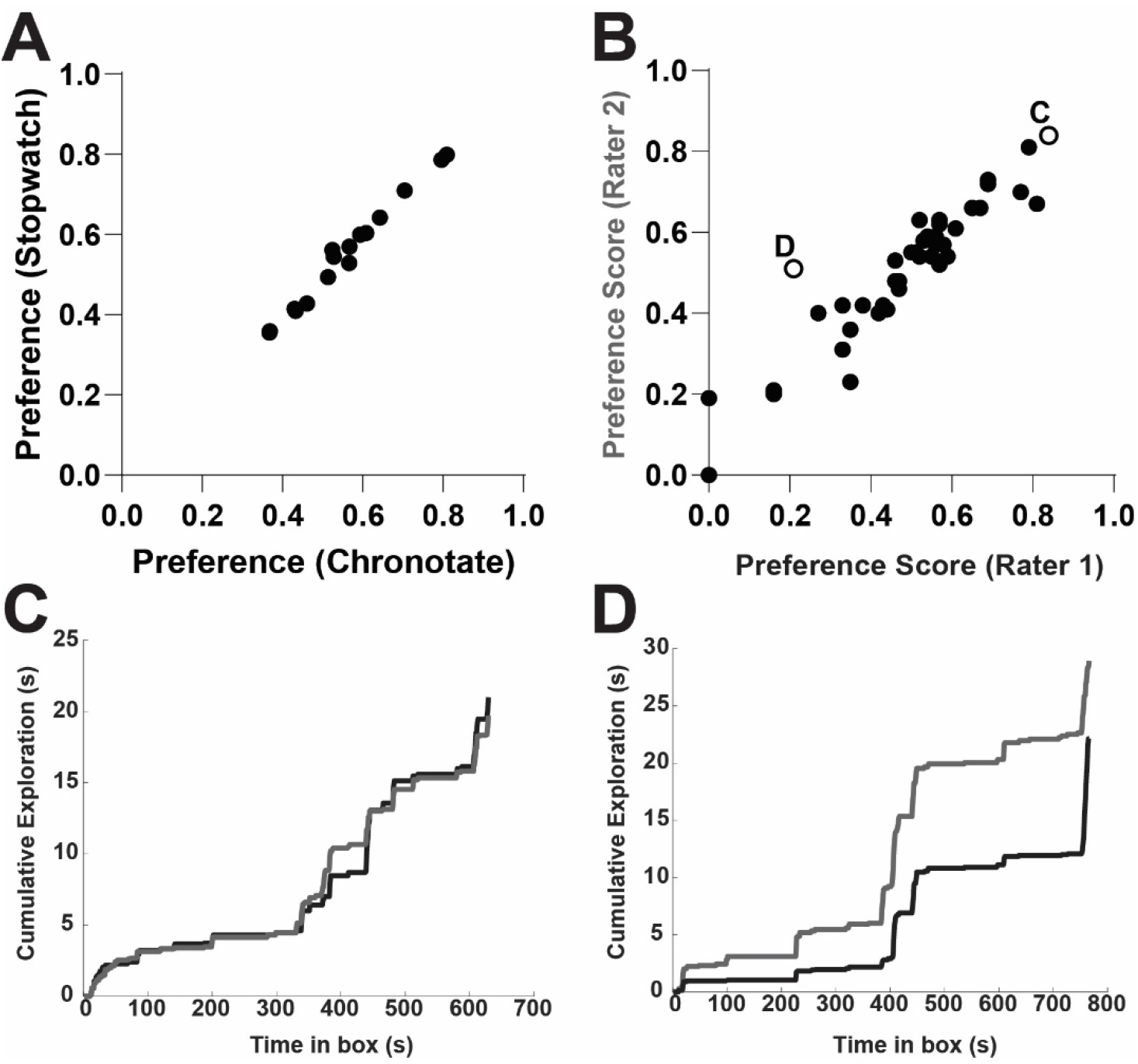
Time-stamped outputs allow detailed assessment of inter-rater reliability. Preference scores from Chronotate are highly correlated to those scored with a stopwatch (Pearson’s r=0.99, *p*<0.0001, n=16 videos). (B) Preference scores by two independent raters are highly correlated (Pearson’s r = 0.93, *p <*0.0001, n=42 animals). (C, D) Cumulative exploration across a video with high inter-rater agreement (C) and a video with lower inter-rater agreement (D). The points corresponding to these two videos are noted in B.

**Figure 3:**
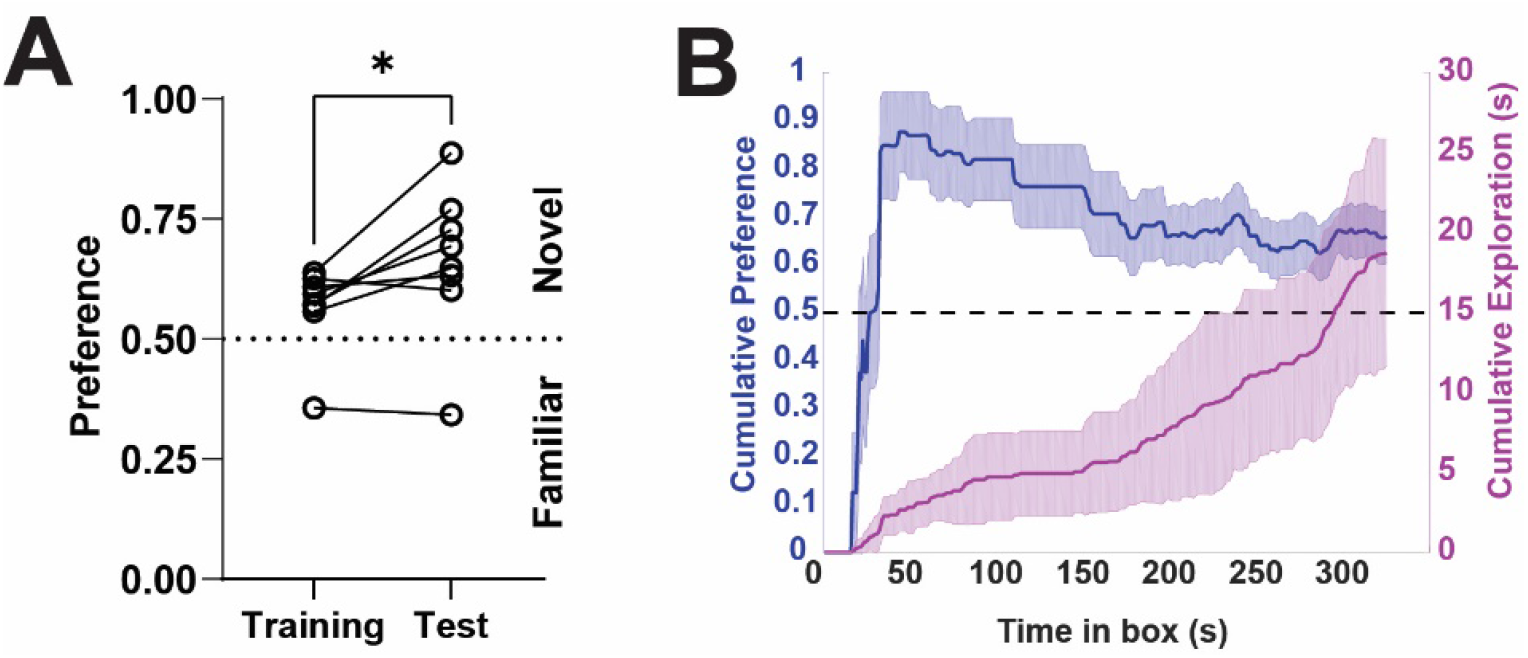
Determining the time-course of object preference across the testing session. (A) Animals successfully learned to discriminate between the novel and familiar object-location pair during the test session (paired t-test *p*=0.028, n=8 animals), with animals showing a preference for the novel object-location. (B) Mean cumulative preference for the novel object over the course of the session (blue, left axis) and mean cumulative total object exploration time (magenta, right axis). Preference for investigating the novel object-location peaks at approximately 45 seconds after the start of the session, which corresponds to an average combined exploration of both objects of 2.7 seconds. Shaded error bars represent SEM.

### Scoring

Mouse-object interactions were scored manually using stopwatches or using Chronotate [12]. Interactions were identified as periods in which an animal was sniffing, whisking at the object, or looking at an object while touching it. Time spent sitting on objects, touching them while rearing up, or grooming while facing an object were not counted as exploration. The mouse’s preference for investigating the moved (novel) vs the unmoved (familiar) objects was calculated as the ratio of time spent interacting with the novel over the total time:

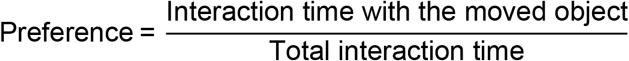

Additionally, for the purpose of excluding animals that showed a strong preference during the training, a discrimination index (DI) was calculated:

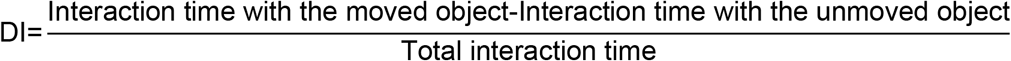

Mice that had a strong preference (DI > 0.25 or < -0.25) for investigating one object over another during the training session or who exclusively interacted with one object during the test session were excluded.

For scoring with stopwatches, total time spent with each object was scored with an individual stopwatch for each of the two objects, starting and stopping the timer at the beginning and end of each bout of exploration [12]. For scoring with Chronotate, each of the two objects was assigned to a particular keyboard key. Each bout of object interaction was marked by pressing the relevant key at the start of exploration and releasing the key at the end of the bout. Marker files containing time stamped interaction bouts for each video were exported for subsequent analysis.

### Chronotate

Chronotate uses key-down and key-up keyboard events to record video timestamps of the start time and end time of each bout of interaction (**Fig 1C**). Simultaneous scoring of up to four different objects is supported. Because the recorded timestamps are relative to the actual position in the video and not an external timer, there is no need to watch the video back in real-time while scoring. Chronotate allows the user to speed up video playback to score long videos more efficiently, or slow down playback to more precisely mark the times of rapid events, all with keyboard strokes (**Fig 1C**). Additionally, Chronotate allows the user to skip forward or backward in fixed increments, and seek to a specific time using a progress bar integrated into the graphical interface (**Fig 1B**).

Each interaction start/stop event logged is displayed in the side panel for review (**Fig 1B**). Following scoring, these markers can be exported and analyzed externally according to the needs of the individual experiment. In addition, a previously exported marker file can be imported alongside its corresponding video in order to review and assess inter-rater reliability. Importantly, clicking on any event recorded in the side panel will seek the video playback to the recorded timestamp, allowing an experimenter to review the scoring of any individual event.

All functionality of Chronotate is implemented in Python. The graphical interface (**Fig 1B**) is implemented using the tkinter package[13]. Video playback is supported by a modified version of the open-source tkVideoPlayer package. The import and export of csv data is implemented using the pandas package[14]. To run the source files, we recommend setting up a new Conda environment for Chronotate with these dependencies.

All source files along with a precompiled executable file (.exe) are available from the Shuman Lab’s GitHub page (https://github.com/ShumanLab/Chronotate). The precompiled .exe is a standalone application compatible with Windows 10 & 11 and requires no installation. The source files require Python 3.6-3.10 and are cross-platform compatible with OSX, Linux and Windows.

### Data Analysis & Statistics

The marker output files from Chronotate were processed by a custom MATLAB script (available on the Shuman Lab’s GitHub) to quantify the total exploration time with the novel and familiar objects. The cumulative exploration with each object was calculated on a second-by-second basis to generate a curve of preference for the novel object over the course of the session. All statistical tests were carried out using GraphPad Prism 9.

## RESULTS

We first show that Chronotate does not introduce a bias into the resulting preference scores. A set of 16 NOL behavior videos were each scored twice by a blinded experimenter, one using stopwatches and a second time using Chronotate. Preference scores were highly correlated (**Fig 2A**: Pearson’s r=0.99, *p*<0.0001, n=16 videos), indicating high agreement between the two scoring methods.

One of the major benefits of using Chronotate to score NOR/NOL video data is the ability to review scoring between raters on a level that is more granular than standard approaches. Typically, inter-rater reliability on these tasks is compared by determining how similar final preference, or discrimination index, values are to each other, as we show in **Fig 2B** (Pearson’s r=0.93, *p*<0.0001, n=42 animals, comparing preference scores between two independent raters). Unfortunately, however, using this method to troubleshoot videos where two raters have low inter-rater reliability presents the challenge of determining where in the video a difference in scoring emerged. Fortunately, because Chronotate outputs each time-stamped exploration bout, the moments where two raters diverged in their scoring becomes easily identifiable (**Fig 2C, 2D**). These portions of the video can then be isolated, reviewed, and re-scored as needed.

The NOR/NOL tasks leverage rodent’s innate tendency to investigate a novel stimulus or location over a familiar one. As such, our mice trained in the NOL task showed a preference for investigating the novel object-location pair that emerged during the test session (**Fig 3A**, paired t-test, *p*=0.028, n=8 animals, novel object-location preference during training 0.57 ± 0.03, preference during test 0.66 ± 0.06). This indicates that the mice had a memory of the familiar object-location pair from the training session and were therefore able to identify which object had moved. However, novelty is a time-limited phenomenon, and as a rodent spends increasing time in the arena on the testing day, the novelty of the moved object will, in theory, begin to diminish. Using Chronotate to log object exploration and preference on a cumulative bout-by-bout basis, we can see that mice show a maximal preference for exploring the novel object-location after 2.7 seconds of total exploration, which occurred 45 seconds into the test session (**Fig 3B**, n=8 animals), and that as the mice begin to spend more time with the objects, this preference is diminished.

## DISCUSSION

Here, we describe Chronotate and provide examples of its utility for facilitating manual scoring of NOR and NOL behavior videos. In short, Chronotate is an open-source graphical user interface implemented in Python that allows the user to score behavior videos with keyboard strokes and outputs timestamped bouts of interest. We show that this time-stamped data can be used to isolate key events, for example, where two raters diverge in their scoring of the data (**Fig 2D**). In addition, this time-stamped data can be used to determine if an object-location preference may have been diminished by the duration of the testing session or the amount of total exploration (**Fig 3**).

An additional benefit of Chronotate is the ability to skip forward through portions of behavior video in which an animal is not in proximity to an object of interest, reducing the amount of time raters must spend reviewing each video. However, a future add-on to this software could integrate open-source location tracking modules, such as the one implemented in ezTrack [9] to automatically direct the user to portions of the video in which the animal is close enough to interact with the object, requiring the rater to review only the timepoints when object interaction is possible.

While we describe Chronotate’s utility for NOR/NOL videos, its potential applications far exceed this scope. Not only can Chronotate be adapted for the scoring of any videos requiring manual, time-stamped notation, but future refinements to automated behavior scoring tools will hopefully reduce the need to perform manual scoring of behavior videos. In this scenario, Chronotate may prove useful for generating labeled training data to be passed into supervised machine learning models.

## ACKNOWLEDGEMENTS

We would like to thank all members of the Shuman and Cai labs for their comments and feedback throughout the implementation of Chronotate. This work was supported by NIH grants R01 NS116357 (TS), RF1 AG072497 (TS), R01 MH120162 (DJC), DP2 MH122399 (DJC), R56 MH132959 (DJC), F32 NS116416 (ZCW), F31 AG069496 (LV), and an AES Predoctoral Fellowship (YF).

## AUTHOR CONTRIBUTIONS

PAP and TS conceptualized Chronotate. PAP designed and implemented Chronotate and performed data analysis. ZCW offered conceptual feedback. NV, KD, AG, AJ, LV, LC, YF designed and performed behavioral testing and video scoring. ZCW, PAP, DJC, and TS wrote and edited the manuscript. DJC and TS supervised the project.

